# Automated Cyclic Super-Resolution Microscopy for Nanoscale Protein Mapping

**DOI:** 10.64898/2026.03.16.712232

**Authors:** Hongqiang Ma, Chaojie Zhang, Shuyuan Zheng, Shengwei Chen, Yang Liu

## Abstract

Nanoscale mapping of multiple molecular targets is essential for decoding cellular architecture, but current approaches are limited by low throughput, intensive manual intervention, and signal variability across imaging cycles. Here, we introduce CycSTORM, an integrated super-resolution imaging platform for automated cyclic (d)STORM that enables multiplexed nanoscale protein mapping in single cells. To overcome the instability inherent in multi-day imaging, CycSTORM combines automated fluidic exchange, active 3D drift correction, and an oxygen-excluded imaging environment to stabilize fluorophore blinking and enables sub-5 nm registration across day-long experiments. Furthermore, CycSTORM incorporates a rapid chemical inactivation step using meta-chloroperoxybenzoic acid (mCPBA), eliminating >99.9% residual fluorescence within 10 minutes and minimizing inter-cycle crosstalk while preserving sample integrity. By standardizing labeling to Alexa Fluor 647, a fluorophore with stable and well-characterized blinking behavior, CycSTORM minimizes fluorophore-dependent variation and provides a robust platform for consistent localization precision across cycles, enabling reliable mapping of protein organization in single cells. Using CycSTORM, we simultaneously map multiple protein targets within the same cells with nanometer precision. Together, these advances transform cyclic super-resolution imaging into a scalable approach for quantitative nanoscale mapping of molecular organization in single cells.

## Introduction

Cellular behavior depends not only on protein composition but also on how proteins are spatially arranged within individual cells. However, directly mapping this molecular organization in situ at nanoscale resolution remains challenging. Spatial proteomic imaging has revealed important spatial relationships among cell types across a large cell population within tissue microenvironment, yet it does not quantitatively compare the nanoscale organization of multiple proteins inside the same cell^1–4^. Single-molecule localization microscopy (SMLM) localizes proteins with nanometer precision^5,6^, and has enabled direct visualization of intracellular molecular organization.

However, most SMLM studies remain limited to only a few targets^7^. Systematic mapping of multiple proteins requires multiplexing, yet current implementations rely heavily on manual intervention and are difficult to scale^8,9^. Simultaneous multi-color SMLM has been explored to increase multiplexing within a single acquisition by spectrally separating fluorophores^10,11^. In practice, spectral multiplexing is constrained by emission overlaps and steric crowding in densely labeled samples. In addition, differences in fluorophore brightness and blinking kinetics lead to target-dependent localization precision, preventing reliable quantitative comparison across proteins^10,11^.

Cyclic SMLM imaging strategies address spectral limitations by sequential labeling, imaging, and signal removal across multiple rounds. DNA-based methods such as DNA-PAINT and its variants achieve high multiplexity through transient hybridization of fluorescent imager strands^12–16^. However, these approaches rely on custom oligonucleotide design and extensive optimization of strand sequences, binding kinetics, target-dependent labeling performance, and imaging conditions, increasing experimental complexity and limiting routine use. In addition, freely diffusing fluorescent imager strands generate substantial background fluorescence, and effective suppression often relies on total internal reflection (TIRF) or highly inclined illumination (HILO), which limits imaging depth in thick samples and reduces fields of view in large samples. Cyclic (d)STORM^7–9,17^, based on conventional antibodies and Alexa fluorophores, provides a more accessible route for multiplexed super-resolution imaging and is compatible with simple widefield configurations.

Despite this progress, reliable multi-cycle super-resolution imaging remains challenging. Residual fluorescence from previous labeling rounds must be removed to prevent inter-cycle crosstalk, and commonly used signal-removal procedures such as antibody elution and photobleaching, can be time-consuming and may affect epitope integrity. Accurate registration of protein targets imaged in different cycles further requires three-dimensional drift correction, typically implemented using fiducial markers with additional sample preparation^9,17^. In addition, the limited stability of imaging buffers over extended acquisition necessitates controlled reagent exchange, increasing experimental complexity^17^.

Here we present CycSTORM, an automated platform for cyclic (d)STORM imaging designed to enable routine multiplexed nanoscale protein mapping in single cells. The system integrates three practical advances that address key barriers to multi-cycle super-resolution imaging. First, an automated staining and imaging workflow within a unified control framework enables sequential labeling and nanometer-scale 3D registration over day-long experiments, while active brightfield registration-based drift correction provides 3D nanometer-scale stability without fiducial markers. Second, rapid chemical fluorophore inactivation using meta-chloroperoxybenzoic acid (mCPBA) removes >99.9% fluorescence residual within 10 minutes, eliminating inter-cycle carryover without prolonged photobleaching or harsh elution. Third, a nitrogen-purged imaging environment during staining and imaging stabilizes redox conditions and preserves consistent single-molecule blinking. By standardizing fluorophore photophysics and minimizing perturbation between labeling rounds, CycSTORM enables scalable mapping of multiple protein targets within the same cells over large imaging areas. This framework paves the way for systematic nanoscale analysis of intracellular protein organization across many cells.

## Results

### CycSTORM platform and automated cyclic workflow

To enable multiplexed spatial mapping of nanoscale molecular organization, we developed CycSTORM, an automated platform for cyclic (d)STORM staining and imaging, as shown in Fig. 1. CycSTORM perform iterative cycles of antibody labeling, super-resolution imaging, and rapid chemical fluorophore inactivation within an automated framework. The platform integrates a high-stability super-resolution localization imaging system (SMLM) coupled with an automated fluidic exchange unit (referred to as AutoStainer). The SMLM system is configured for wide-field (d)STORM imaging and high-speed image acquisition at a large field of view (100×100 μm^2^)^18^. One key component of the automated CycSTORM system is the real-time 3D drift correction system based on bright-field imaging of the diffraction pattern from the inherent cell or tissue architecture, enabling drift-correction accuracy for sub-2 nm lateral and sub-5 nm axial dimensions.

**Figure 1.**
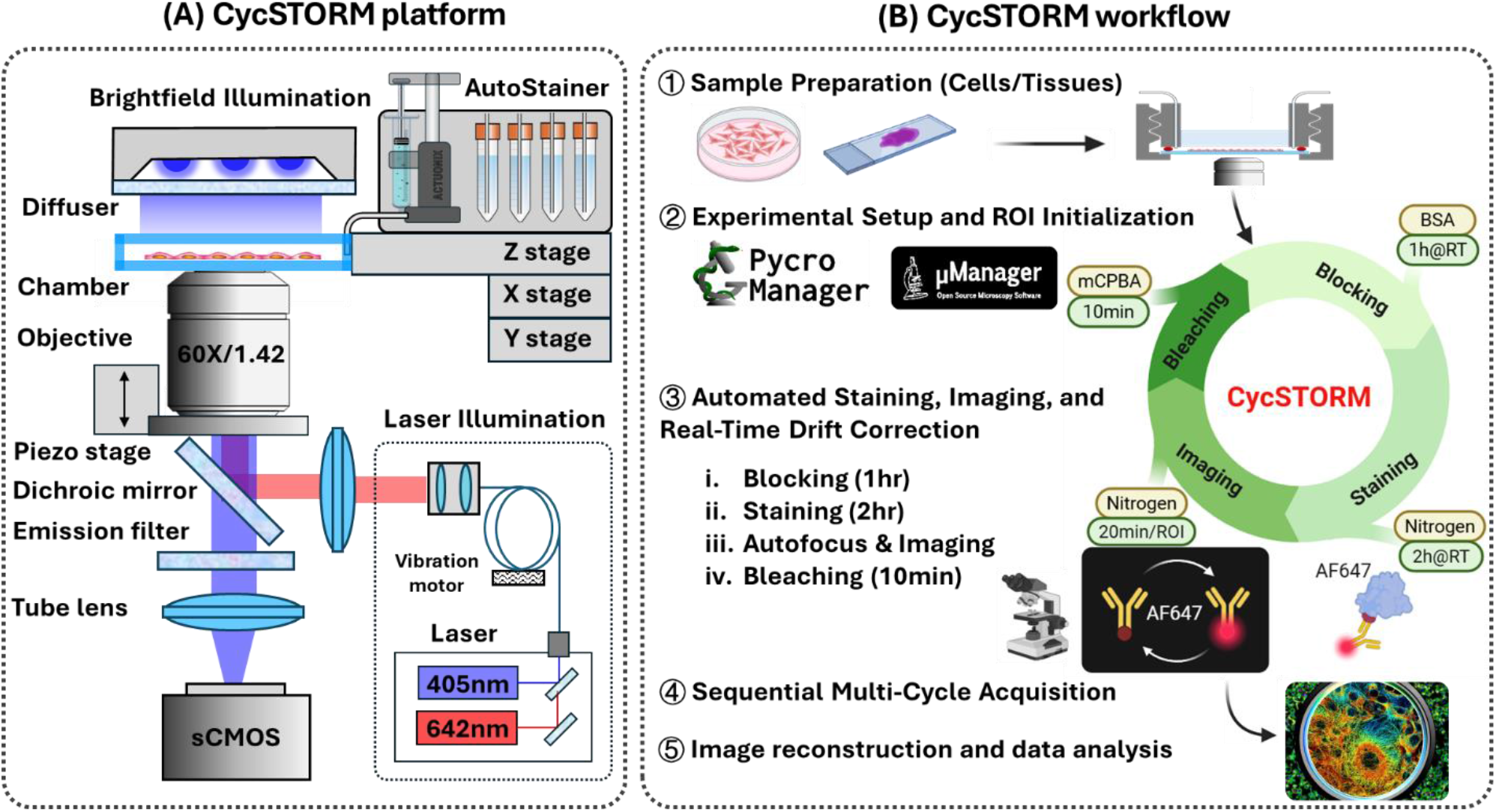
CycSTORM automated platform and cyclic multiplexing workflow. (A) Integrated imaging-fluidics setup. A custom widefield microscope equipped with a 60×/1.42 NA objective, 642 nm excitation (405 nm activation), and an sCMOS camera is coupled to a motorized XYZ stage and high-speed Z-piezo for active nanometer-scale drift correction. Automated reagent exchange is performed by the AutoStainer module using a syringe pump and multiport valves to deliver antibodies, imaging buffers, and fluorophore inactivation agents. (B) CycSTORM workflow. System control and synchronization are implemented through Python-based software. Each cycle consists of antibody labeling, automated washing, autofocusing, dSTORM acquisition, online drift correction, and rapid fluorophore inactivation with mCPBA, enabling repeated imaging rounds with minimal inter-cycle carryover.

The CycSTORM workflow converts manual super-resolution imaging into an automated imaging-staining pipeline. Samples mounted in a custom-designed flow chamber directly coupled to the AutoStainer. The AutoStainer (Fig. 4A) employs a precision syringe pump and multiport switching valves to deliver blocking buffer, antibody solutions, imaging buffer, wash solutions, and fluorophore inactivation agents with microliter-scale control. All hardware components, including laser excitation, stage motion, camera acquisition, autofocusing, drift correction and fluidic exhange, are automated through a Pycro-Manager^19^-assisted Python-based control framework (Fig. 6), enabling fully autonomous multi-day operation without user intervention.

Each cycle consists of four major steps: blocking, antibody incubation, (d)STORM imaging, and fluorophore inactivation. Antibody incubation is performed using AF647-conjugated primary or secondary antibodies, with standardized labeling conditions across all rounds. By constraining each cycle to the optimized photophysics of AF647, CycSTORM maintains consistent blinking kinetics, photon yield, and localization precision across targets.

A key advancement of CycSTORM is rapid chemical fluorophore inactivation using meta-chloroperoxybenzoic acid (mCPBA)^20^, which efficiently oxidizes AF647, removing residual fluorescence after each imaging round without producing bubbles. The bleaching reagent is highly effective and achieves >99.9% fluorescence suppression within 10 minutes. After fluorophore inactivation, a brief wash with PBS restores neutral pH, and the system automatically initiates the next staining round.

Throughout repeated cycles, precise spatial registration is maintained through continuous active drift correction and controlled fluidic exchange. This automated sequence of labeling, imaging, and fluorophore inactivation enables dense multiplexing of protein targets while preserving nanoscale spatial fidelity during multi-day experiments.

### Automated 3D Nanometer Image Registration and Drift Correction System

Maintaining nanoscale stability across repeated fluidic exchanges and day-long imaging sessions is a central requirement to CycSTORM. The automated cyclic workflow involves multiple rounds of staining, washing, and imaging, during which mechanical and thermal fluctuations can accumulate and produce sample drift of several microns. Conventional drift correction often relies on fluorescent fiducial markers that may lie in a focal plane different from the sample, requiring alternating focus between the fiducial and the imaging plane during acquisition^9,17,21^. To address this challenge, CycSTORM incorporates an autonomous 3D stabilization pipeline that continuously tracks sample displacement using intrinsic morphological features of the sample without fiducial markers^18,22,23^.

The stabilization framework employs a hierarchical marker-free drift correction strategy that compensates for both large-scale displacements accumulated between imaging cycles and small-scale drift during each STORM acquisition (Fig. 2). At the beginning of the cyclic STORM imaging experiment, a 3D bright-field reference stack is acquired at each imaging area along the axial direction (Fig. 2A➀). This reference stack establishes a structural coordinate for the sample and provides sufficient axial range to capture micro-scale displacement arising from fluidic exchange or long-term thermal drift during multi-cycle staining and imaging. Furthermore, we take advantage the bright-field image stack to reconstruct quantitative phase images (QPI)^24^, which provides intrinsic structural contrast of cellular architecture, from overall cell morphology to fine subcellular features, thereby adding complementary structural information to fluorescence measurements and facilitating detection of morphological deformation between cycles^25^.

**Figure 2.**
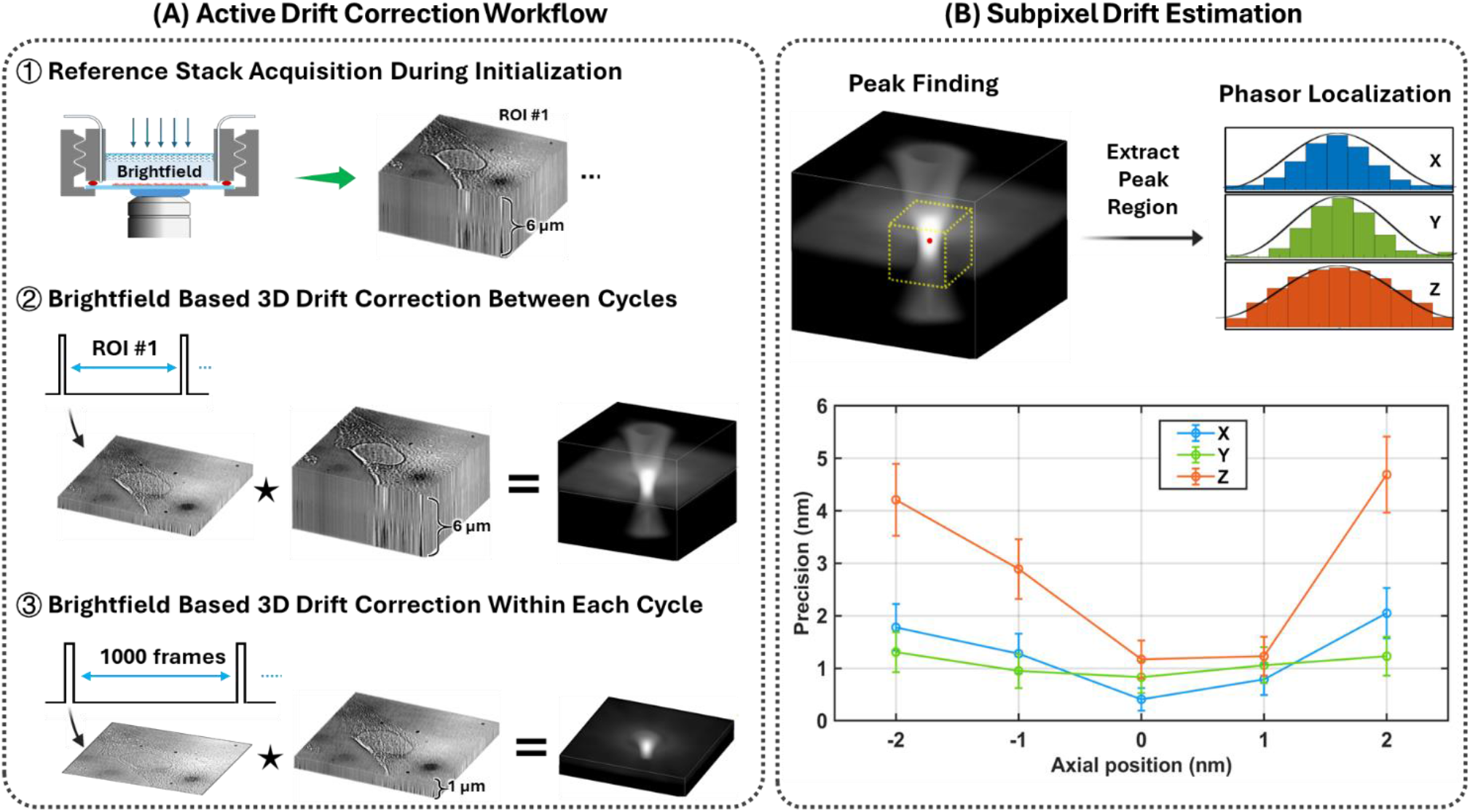
Automated 3D image registration and nanometer-scale drift correction during cyclic imaging. (**A**) Hierarchical marker-free drift correction workflow used in CycSTORM. System stability is maintained through a three-stage stabilization pipeline: (i) acquisition of a 3D bright-field reference stack at each imaging area, (ii) large-range 3D drift correction (∼6 μm) prior to each STORM imaging cycle, and (iii) short-interval 3D drift correction (∼1 μm) during STORM acquisition at each imaging cycle. (**B**) Principle and performance of subpixel 3D drift correction. Pixel-level displacement is first determined from the peak of the 3D normalized cross-correlation volume between the target image and the reference stack. The peak region (11 × 11 × 11 pixels) of the correlation is then refined using a phasor-based algorithm to estimate subpixel displacement X, Y and Z without iterative fitting. This approach enables high-speed, high-precision drift estimation. Residual drift after correction remains below ∼2 nm laterally and ∼5 nm axially over an axial range of ±2 μm, enabling nanoscale system stability during day-long multi-cycle imaging and staining.

Before each imaging cycle following fluorescence staining, a bright-field image stack (100 nm spacing over a ∼1 µm axial range) is acquired (Fig. 2A➁). This stack is used both to quantify 3D morphological deformation and to register the sample to the reference stack. Cross correlation is first used to estimate the accumulated 3D displacement at the pixel level, followed by a refinement step to achieve the subpixel precision using a fast phasor-based localization algorithm^26,27^. This strategy enables rapid computation of the 3D drift vector while maintaining nanometer accuracy.

During STORM acquisition, CycSTORM performs short-interval active drift correction approximately every 30 seconds for nanometer-scale drift occurring during each 10-20 minutes image acquisition. Because intra-cycle drift is typically limited to tens to hundreds of nanometers, the registration search is restricted to a small axial range (Fig. 2A➂), enabling fast computation while maintaining high precision.

Together, these hierarchical correction steps enable stable nanoscale stability across extended imaging sessions. As shown in Fig. 2B, the system maintains sub-nanometer lateral precision and <2 nm axial precision at the focal plane, remaining below 5 nm even with ∼2 μm defocus. After SMLM localization and reconstruction, residual drift is further corrected in the localization dataset every 100 frames (∼3 s) using Adaptive Intersection Maximization (AIM) algorithm^28^. This final refinement removes high-frequency drift components and ensures stable nanoscale registration throughout day-long cyclic imaging experiments. Details of the registration algorithm are described in the Materials and Methods.

### Quantitative Evaluation of Fluorophore Inactivation Efficiency for CycSTORM

Reliable cyclic super-resolution imaging requires efficient removal of residual fluorescence from previous labeling rounds to prevent inter-cycle crosstalk. To evaluate fluorophore inactivation in CycSTORM, we examined cells labeled with Alexa Fluor 647 (AF647)-conjugated antibodies against histone H3 lysine 9 trimethylation (H3K9me3) (Fig. 3A). H3K9me3 forms densely packed heterochromatin domains within the nucleus, allowing residual fluorescence after fluorophore inactivation to be readily detected.

**Figure 3.**
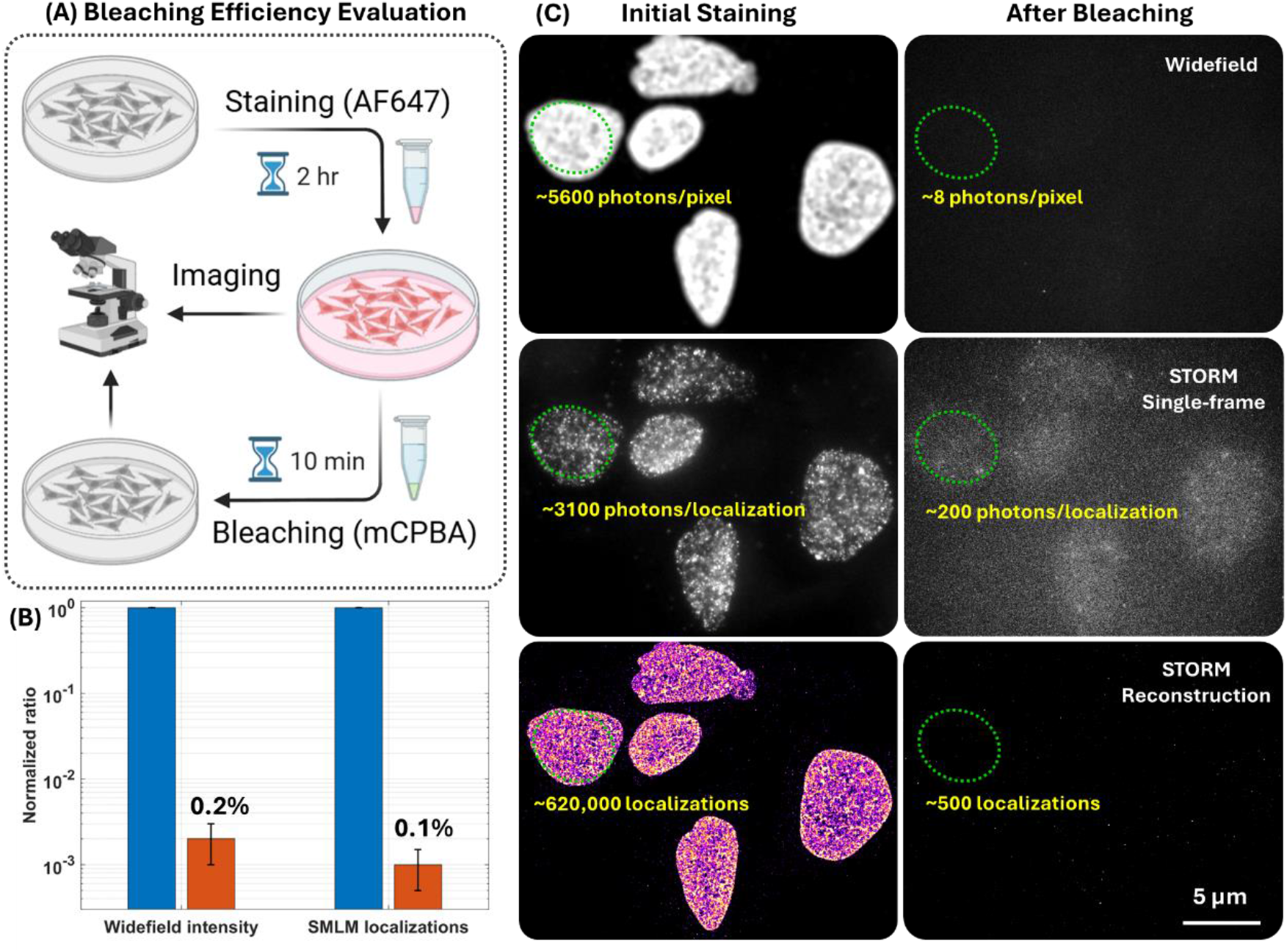
Quantitative evaluation of fluorophore inactivation efficiency for cycSTORM. (**A**) Workflow for assessing fluorophore inactivation efficiency: cells were initially stained with H3K9me3-AF647 (2 hour, room temperature), followed by conventional widefield imaging (50 W/cm^2^) and dSTORM imaging. A 10-minute mCPBA fluorophore inactivation step was applied, after which widefield and dSTORM imaging were repeated to assess residual signal. (**B**) Comparison of the AF647 intensity in the wide-field image and the number of localizations in the reconstructed STORM images before and after mCPBA-based fluorophore inactivation. (**C**) Representative images showing widefield (∼5600 photons/pixel vs. 8 photons/pixel), single-frame raw image of dSTORM (∼3100 photons/localization vs. ∼200 photons/localization), and reconstructed dSTORM (∼620,000 localizations vs. 500 localizations in the circled regions) before and after mCPBA-based fluorophore inactivation.

**Figure 4.**
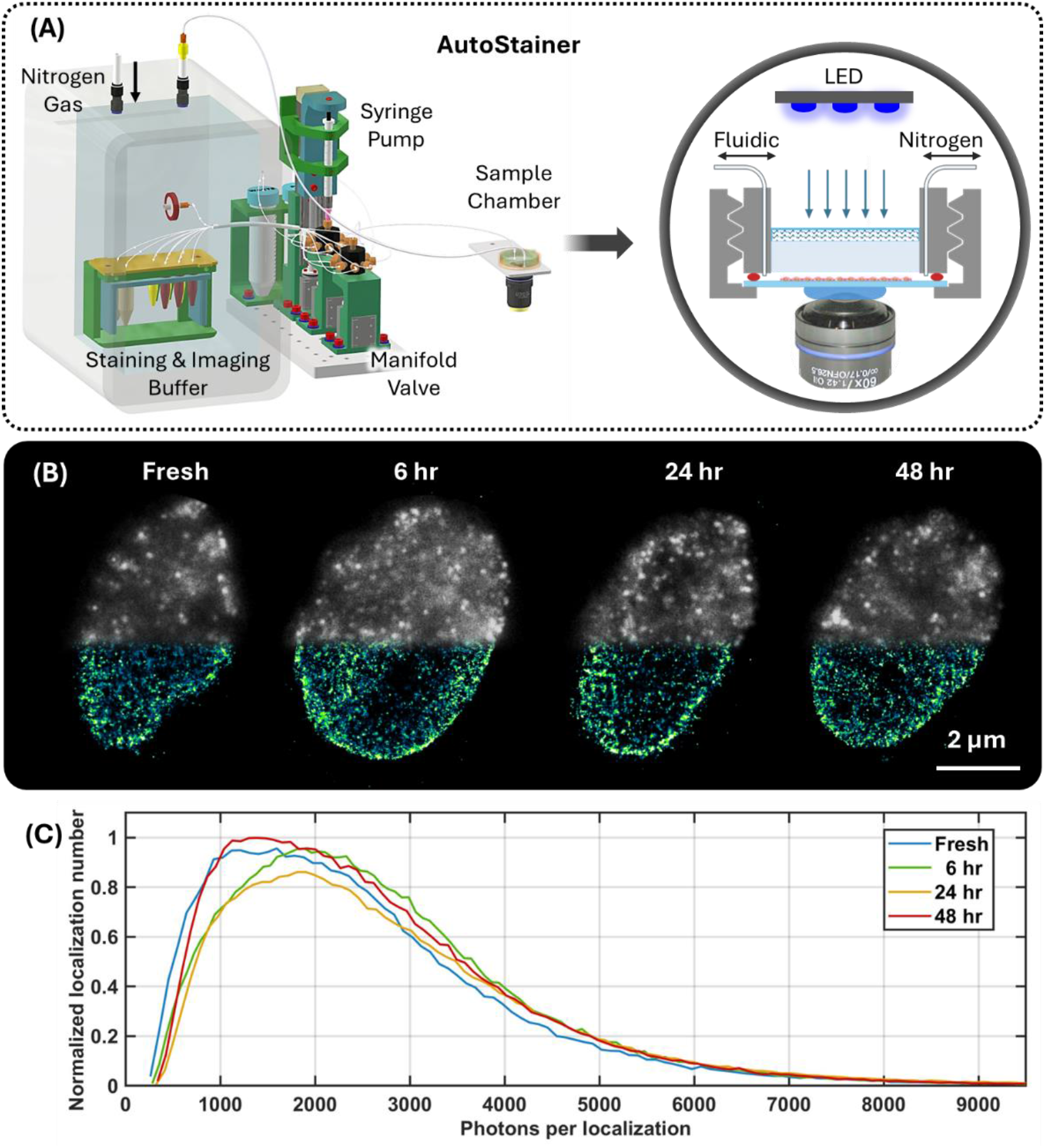
Nitrogen-purged AutoStainer system enables stable long-term dSTORM imaging. (**A**) Schematic of the AutoStainer integrated with a nitrogen-purged sample chamber for automated delivery of staining reagents and imaging buffer. The system includes a syringe pump, multiport valves, and imaging and staining buffers maintained under nitrogen environment to prevent oxygen-induced degradation of fluorophore blinking. The inset illustrates the sample chamber with the fluidic delivery of nitrogen-purged staining or imaging buffers. (**B**) Temporal stability of dSTORM imaging. Representative raw single-frame blinking events (top half) and reconstructed super-resolution images (bottom half) are shown in the presence of imaging buffer stored in the nitrogen-purged chamber for 0 (fresh), 6, 24, and 48 hours. (**C**) Quantitative assessment of fluorophore photoswitching performance. Distributions of normalized localization numbers versus photons per localization remain stable over 48 hours, indicating preserved fluorophore blinking efficiency and consistent localization precision under nitrogen protection.

Cells were first imaged to establish a pre-bleach state. As shown in Fig. 3C, conventional widefield fluorescence imaging under moderate illumination confirmed nuclear labeling and strong fluorescence signal, and the subsequent dSTORM imaging resolved dense heterochromatin nanodomains characteristic of H3K9me3. Samples were then treated with 10 mM meta-chloroperoxybenzoic acid (mCPBA) for 10 minutes, delivered via the automated fluidic system described above. After washing with PBS, widefield and STORM imaging were repeated under identical settings to directly quantify residual fluorescence (Fig. 3A).

As shown in Figs. 3(B-C), quantitative analysis revealed a dramatic decrease in fluorescence signals after mCPBA treatment. Conventional widefield fluorescence intensity of H3K9me3 decreased from ∼5,000 photons (100% baseline) to fewer than 8 photons (<0.2%) per pixel, approaching the detection noise floor. Analysis of the raw dSTORM frame further showed the average photon count per localized individual single molecules decreased from ∼3100 photons to 200 photons.

Single-molecule localization analysis provided an even more stringent assessment. The total number of localized molecules in the nucleus associated with pre-bleached heterochromatin clusters were reduced to less than 0.1% of their original value (>99.9% reduction). Representative images further illustrate the effect of mCPBA bleaching (Fig. 3C), where strong pre-bleach fluorescence and dense STORM localizations (∼620,000 localizations within the selected region) were nearly completely eliminated after treatment (∼500 residual localizations). The few remaining localizations were sparsely distributed and showed no spatial correlation with the original chromatin architecture, confirming effective elimination of structured fluorescence signal.

Together, these results demonstrate that mCPBA achieves near-complete AF647 inactivation within just 10 minutes. The suppression of bulk fluorescence and stochastic blinking ensures negligible inter-cycle crosstalk, allowing immediate re-staining of the next imaging target without detectable interference from prior imaging rounds. Combined with automated fluidic delivery, this rapid chemical inactivation provides a robust foundation for highly multiplexed cyclic super-resolution imaging with AF647-labeled targets.

### Nitrogen-purged Automated Fluidic System for Long-term Cyclic dSTORM Imaging

A critical challenge in long-term cyclic dSTORM imaging is the limited lifetime of the imaging buffer. The oxygen-scavenging components required for efficient fluorophore blinking gradually degrade over time, leading to reduced blinking efficiency and localization precision^29^. To maintain stable imaging conditions throughout extended cyclic experiments, we developed a custom-built automated staining and buffer-exchange system (AutoStainer) integrated with a nitrogen-purged environmental chamber^30^.

As illustrated in Fig. 4A, the AutoStainer consists of a precision syringe pump and a multiport valve that deliver staining reagents and imaging buffers into a sealed sample chamber mounted on the microscope stage. The chamber is continuously purged with nitrogen, creating an oxygen-minimized environment that preserves the stability of the dSTORM imaging buffer during extended imaging sessions while minimizing mechanical disturbance during reagent exchange.

To evaluate the temporal stability of this nitrogen-purged system, we monitored dSTORM blinking performance over a 48-hour period using the imaging buffer stored in the chamber. We compared the raw single-frame blinking events and the reconstructed super-resolution images using imaging buffers stored for 0 (fresh), 6, 24, and 48 hours in the nitrogen-purged chamber. As shown in Fig. 4B, the localization density and reconstructed chromatin structure of H3K9me3 remain visually similar over two days.

To quantitatively assess the effect of the nitrogen-purged environment on the fluorophore photophysics, we analyzed the distribution of photons per localization, a key determinant of theoretical localization precision, thus reconstructed image resolution. The distribution of the total photons per localization remained stable throughout the testing period of 48 hours, with the peak of the photon distribution consistently maintained between approximately 1,500 to 2,000 photons. This consistency ensures that the achievable resolution does not deteriorate as the experiment progresses. In addition, the total number of detected localizations remained high, indicating that the nitrogen purging successfully preserved the oxygen-minimized environment required for efficient fluorophore blinking over time. Together, these results demonstrate that the nitrogen-purged AutoStainer system maintains stable dSTORM imaging conditions for at least 48 hours. This stability is crucial for CycSTORM, where multi-cycle imaging experiments spanning six or more cycles can extend a day or longer and require stable imaging buffer that provides reproducible photo-switching performance from the first cycle to the last.

### Quantitative Phase and Six-Plex Cyclic STORM Imaging of Cellular Architecture and Epigenetic States

To demonstrate both the multiplexed nanoscale imaging capability of CycSTORM, we performed a six-plex cyclic STORM experiment in mouse fibroblast cells (NIH3T3), targeting structural and epigenetic features within the same cell (Fig. 5). Conventional multi-color STORM is typically restricted to two or three targets due to spectral overlap among compatible fluorophores^31,32^. In contrast, CycSTORM expands multiplexing capability through iterative cycles of labeling, imaging, and fluorophore inactivation. The automated workflow described above enables sequential imaging cycles without manual intervention while maintaining stable imaging conditions and nanometer-scale spatial registration across extended experiments. As a result, protein targets imaged in different cycles can be directly overlaid with high spatial fidelity, allowing quantitative comparison of intracellular structures across labeling rounds.

**Figure 5.**
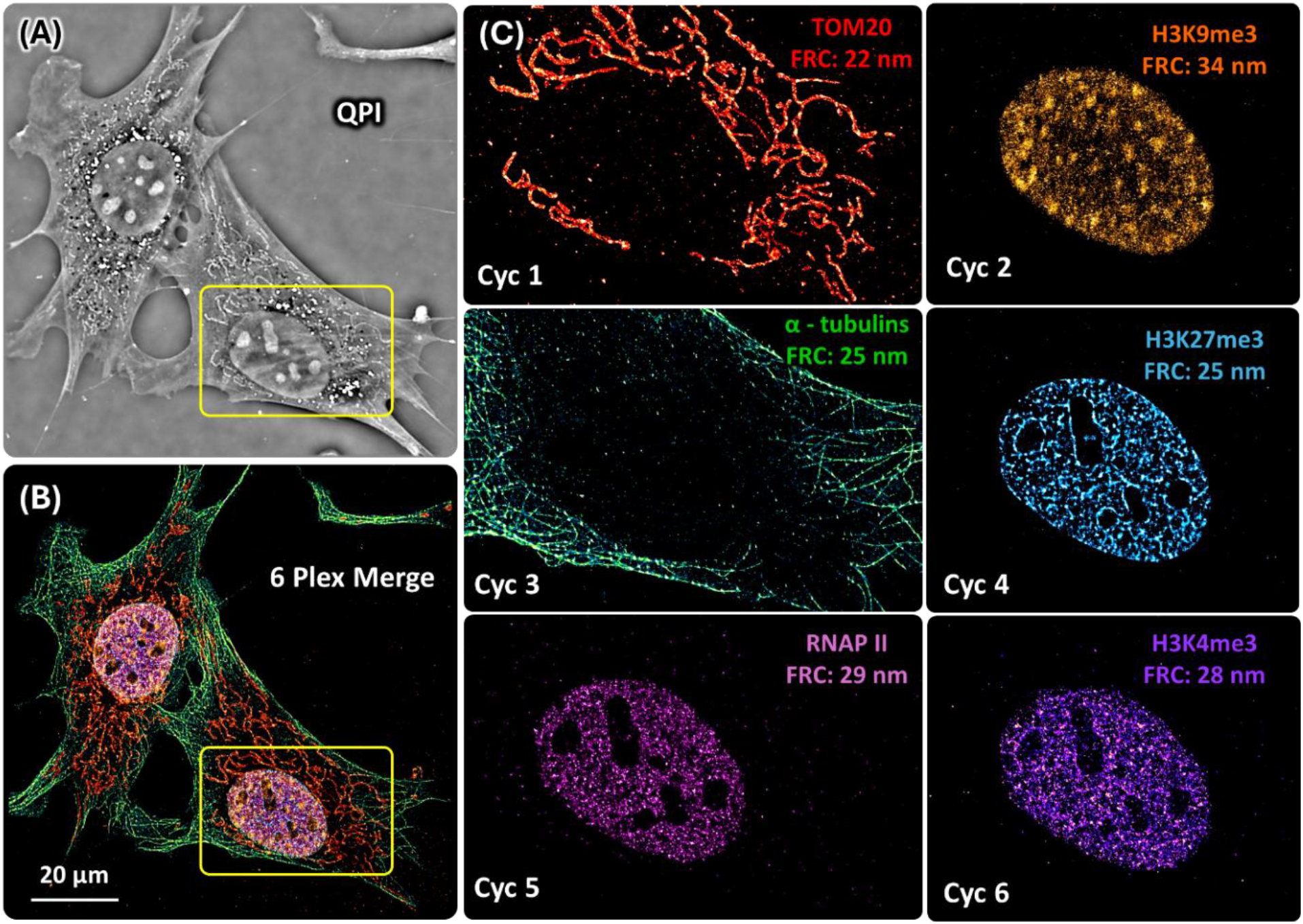
Six-plex cyclic STORM imaging of NIH3T3 cells. (A) Reconstructed quantitative phase image (QPI) generated from the bright-field image stack. (B) Color-merged STORM super-resolution image of six protein markers, suggesting precise spatial registration maintained across all imaging cycles. Scale bar: 20 μm. (C) Super-resolution images of each of the six targets. This panel shows the mitochondrial outer membrane (TOM20), the microtubule cytoskeleton (α-tubulin), and four nuclear epigenetic markers (H3K9me3, H3K27me3, H3K4me3, and RNAPII). This multi-round experiment demonstrates the ability of cyclic STORM to resolve six distinct molecular targets within the same cell.

**Figure 6.**
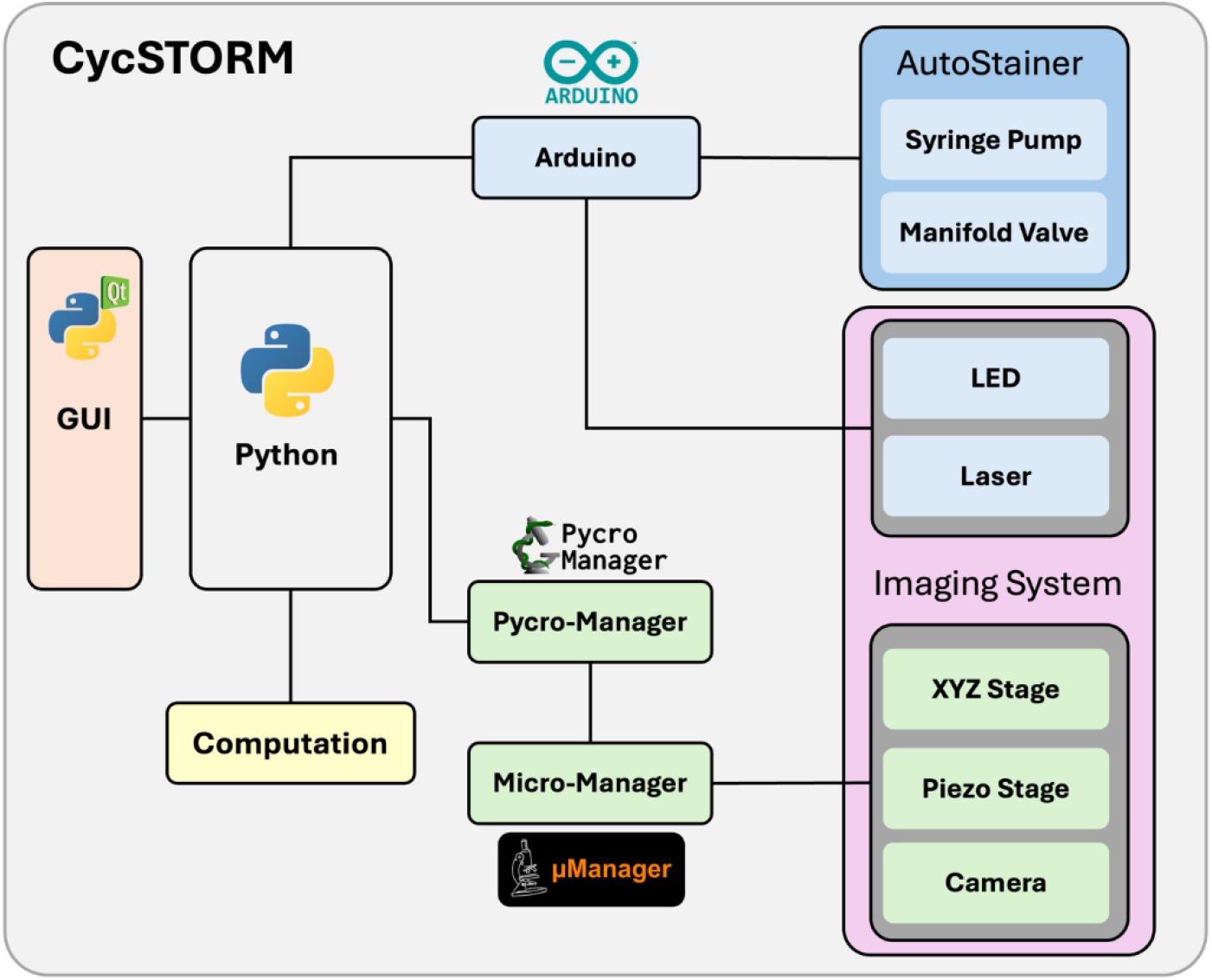
Software and hardware control of the CycSTORM platform. The CycSTORM graphical user interface (GUI), implemented in Python, coordinates the control of light source, motion controls, image acquisition and fluidic exchange for automated cyclic STORM imaging.

Using this approach, we sequentially imaged six targets within the same cells, including the mitochondrial outer membrane marker TOM20, the microtubule cytoskeleton marker α-tubulin, and four nuclear epigenetic markers, H3K9me3, H3K27me3, and H3K4me3, and RNAPII, all on the same cell^24,33^. These targets span cytoplasmic structural networks and nuclear epigenetic states, allowing direct interrogation of spatial relationships between cellular architecture and chromatin organization.

The reconstructed QPI image (Fig. 5A) reveals rich structural information across the same field of view, including fine filamentous features that correspond to the α-tubulin labeling, mitochondrial-associated contrast that colocalizes with TOM20 labeling, and overall nuclear architecture. These results highlight the ability of QPI to provide complementary structural context alongside fluorescence imaging.

Super-resolution imaging revealed distinct nanoscale features for each target (Fig. 5C). Across all targets, reconstructed images achieved Fourier Ring Correlation (FRC) resolutions^34^ between 22 and 34 nm. TOM20 labeling resolved a dense and highly branched mitochondrial outer membrane network distributed throughout the cytoplasm. α-Tubulin imaging delineated microtubule filaments extending toward the cell periphery. Within the nucleus, chromatin organization exhibited characteristic spatial patterns of four epigenetic marks. H3K9me3 occupies highly compact large heterochromatin nanodomains, whereas H3K27me3 formed broader repressive chromatin regions. In contrast, H3K4me3 and RNA polymerase II (RNAPII) with smaller nanodomains were distributed in more dispersed nuclear regions associated with transcriptionally active chromatin. These epigenetic marks displayed distinct spatial distributions and their nanodomains show limited spatial overlap, suggesting the functional compartmentalization of individual epigenetic marks.

The integrity of the six-cycle experiment is evident in the final merged reconstructed STORM image. Despite repeated rounds of staining, imaging, fluorophore inactivation and fluidic exchange, spatial registration based on their intrinsic structural features from the bright-field images remained precise across all targets. The structural features of mitochondrial and microtubule networks are preserved, and the spatial distributions of epigenetic marks were resolved at nanoscale resolution, consistent with the previous reports^35,36^. Importantly, super-resolution performance remained consistent throughout the experiment, with FRC resolutions ranging from 22 to 34 nm across all cycles. These results support that iterative fluorophore inactivation and re-labeling do not compromise ultrastructural preservation or localization precision in cyclic STORM imaging.

## Discussion

Mapping the nanoscale organization of proteins at single-cell level is essential for understanding cellular function. However, achieving highly multiplexed imaging with single-molecule localization microscopy has remained technically challenging, due to its labor-intensive nature and limitations in fluorophore compatibility and spectral crosstalk. In this work, we developed CycSTORM, an automated cyclic super-resolution imaging platform that integrates rapid fluorophore inactivation, a stabilizing imaging environment, nanometer-scale image registration based on intrinsic structural features and reconstruction of high-resolution quantitative phase image from the bright-field reference stack. Together, these components enable multi-cycle imaging with minimal inter-cycle crosstalk while maintaining nanoscale spatial resolution and providing complementary structural information on the same cell.

Several technical advances contribute to the robustness of CycSTORM. First, rapid fluorophore inactivation using mCPBA provides near-complete quenching of Alexa Fluor 647 fluorescence within minutes, effectively eliminating residual fluorescence between imaging cycles. Quantitative measurements demonstrate >99.9% suppression of both bulk fluorescence and single-molecule localizations, ensuring negligible inter-cycle crosstalk during sequential imaging. Second, the nitrogen-purged environmental chamber preserves the efficiency of dSTORM imaging buffer by maintaining oxygen-minimized environment, enabling consistent fluorophore blinking over extended imaging sessions. This stability is essential for cyclic experiments that span many hours or a few days. Third, a hierarchical drift correction strategy based on intrinsic bright-field structural features enables nanometer-scale drift correction and spatial registration without the need for fiducial markers. Importantly, the same bright-field image stack can be used to reconstruct QPI, providing intrinsic structural contrast of cellular architecture across the same imaging area. In our experiments, these high-resolution phase images provided complementary morphological context, revealing fine filamentous features corresponding to the cytoskeleton, mitochondrial-associated structural contrast, and overall nuclear architecture. These capabilities simplify the drift correction workflow and improve confidence in image registration across repeated staining and imaging cycles.

CycSTORM complements existing multiplex super-resolution imaging strategies while emphasizing experimental simplicity and accessibility. DNA-PAINT-based approaches can achieve very high multiplexity through transient DNA hybridization, often exceeding 20 targets^12,13^. However, these methods require custom oligonucleotides conjugated with antibodies, and careful optimization of the binding kinetics of strand sequences, which increases cost and experimental complexity. A further practical challenge is that multiplex imaging across many targets requires careful tuning of imager strand concentrations to match protein abundance, which becomes increasingly difficult when targets span a wide dynamic range of expression levels. In contrast, CycSTORM uses conventional antibodies and a single well-characterized fluorophore (Alexa Fluor 647), eliminating the need for custom conjugation and simplifying experimental preparation while maintaining robust single-molecule localization performance.

The addition of QPI further broadens the information content of each experiment by providing complementary structural context without requiring extra landmark fluorescent labels or additional sample-processing steps.

Other multiplex STORM approaches rely on spectral separation or sequential staining with antibody elution combined with H_2_O_2_-based photobleaching. Spectral STORM^10,11^ expands multiplexity by introducing additional optical elements to distinguish fluorophores based on emission spectra. However, these configurations increase the complexity of the optical setup and can reduce photon collection efficiency, potentially lowering localization precision. In addition, even small levels of spectral crosstalk may complicate quantitative analysis when multiple targets are imaged simultaneously.

Sequential STORM methods have also been demonstrated^7–9,17^, including automated implementations^16,17^, but commonly rely on antibody elution combined with prolonged photobleaching to remove residual fluorescence between cycles. These procedures can expose samples to harsh conditions that may damage epitopes. Moreover, incomplete bleaching can lead to residual fluorescence that accumulates across cycles and becomes detectable after multiple rounds of imaging. In contrast, the rapid fluorophore inactivation strategy used in CycSTORM provides efficient fluorophore quenching within minutes while preserving sample structural integrity. This approach enables repeated imaging cycles with minimal carryover signal, supporting reliable multiplex super-resolution imaging without introducing substantial experimental complexity. In addition, because QPI is reconstructed from the bright-field reference acquisition already required for drift correction, it provides a built-in structural readout that can be used to monitor morphological stability throughout the cyclic workflow.

Despite these advantages, limitations remain. Structural perturbations are inherent to any cyclic imaging approach, as repeated rounds of staining, fluorophore inactivation, and fluidic exchange can introduce small cumulative deformations in biological samples^25^. Although such effects were minimal in the experiments presented here, the maximum number of achievable imaging cycles may ultimately be constrained by the stability of the specimen, which can vary across cell types, substrate coating conditions and tissue preparation. In addition, dye-conjugated primary antibodies are often preferred to minimize IgG cross-binding when multiple targets are labeled sequentially. Future extensions could integrate additional fluorophores, orthogonal labeling strategies, or adaptive imaging workflow to further expand multiplexity and accommodate more complex biological samples. It will also be interesting to explore broader use of the QPI channel, not only for registration or deformation monitoring, but also for joint structural-molecular analysis that links nanoscale protein organization to whole-cell morphology and subcellular architecture.

In summary, CycSTORM provides a practical and scalable framework for multiplex super-resolution imaging. By integrating rapid fluorophore inactivation using mCPBA, a nitrogen-purged imaging environment that preserves the stability of the dSTORM imaging buffer, automated fluidic control, and hierarchical drift correction based on intrinsic bright-field structural features, the platform enables reproducible cyclic imaging while maintaining localization precision and spatial registration across multiple labeling rounds. The six-plex imaging experiments presented here demonstrate that cytoskeletal architecture, mitochondrial structure and nuclear epigenetic states can be sequentially mapped within the same cell with nanoscale resolution, while QPI provides complementary structural context spanning fine subcellular features to overall cellular and nuclear morphology. Because CycSTORM relies on conventional antibodies, a single well-characterized fluorophore, and standard widefield STORM instrumentation, it lowers the experimental barrier for multiplex super-resolution imaging. This combination of simplicity, stability, and automation makes CycSTORM a broadly accessible approach for investigating the spatial organization of protein networks and other molecular assemblies in situ.

## Materials and Methods

### STORM imaging System

The optical system has been described previously^18^. In brief, the super-resolution imaging system is built on a customized inverted fluorescence microscope equipped with an oil-immersion objective lens (UPLXAPO60XO, NA = 1.42, Olympus). A blue LED (425 nm, 3 W), passing through a diffuser, is used as the light source for bright-field illumination. Fluorescence emission is detected using a scientific CMOS camera (ORCA-Flash4.0 V2, Hamamatsu). For STORM acquisition, the central 1024 × 1024 pixel region of the camera sensor was used to ensure imaging within the uniformly illuminated region of the excitation field. Sample positioning and focusing are achieved using motorized XYZ actuators (Z295B, Thorlabs) for coarse lateral and axial positioning, and a piezo-driven nano-Z stage (Mad City Labs, Nano-F100S) for high-precision axial control. Excitation illumination is provided by multiple laser sources including 405 nm (DL405-050, 50 mW, CrystaLaser), 488 nm (DL488-150, 150 mW, CrystaLaser), 561 nm (2RU-VFL-P-2000-560-B1R, 2000 mW, MPB Communications), and 642 nm laser (VFL-P-1000-642-OEM3, 1000 mW, MPB Communications) coupled into a multimode optical fiber (Thorlabs, 400µm core) to generate uniform sample illumination.

### Software and hardware control of CycSTORM platform

The CycSTORM platform integrates a fluorescence microscope, an automated staining module, and a custom Python-based control interface to enable fully automated cyclic STORM imaging including drift correction, image acquisition, immunofluorescence staining and buffer exchange. The overall software and hardware architecture of the system is illustrated in Fig. 6. Microscope hardware components, including the sCMOS camera, motorized XYZ stage, and piezo nanopositioning objective holder, are controlled through Micro-Manager. The CycSTORM control software communicates with Micro-Manager via Pycro-Manager^19^, which allows programmatic control of microscope devices from Python. This software enables automated image acquisition, stage positioning, and focus adjustment during the imaging workflow.

Peripheral devices are controlled through multiple Arduino microcontrollers via serial communication. These devices include LED illumination for bright-field imaging, a laser shutter for switching laser illumination on and off, a servo motor used to adjust laser excitation power through an optical attenuator, and a vibration motor used to reduce speckle artifacts in the laser illumination.

The AutoStainer module is also controlled by an Arduino microcontroller, which performs automated reagent delivery and fluidic operations during staining and bleaching steps. Through this control architecture, the CycSTORM platform synchronizes microscope imaging, reagent exchange, and auxiliary hardware operations, enabling fully automated cyclic STORM imaging.

### 3D Drift Correction and Image Registration

Three-dimensional drift correction was performed using bright-field image registration. At the beginning of each experiment, a reference bright-field image stack is acquired in each imaging area. Images were collected along the axial direction with a step size of 100 nm over a range of 6 μm (user-adjustable). This reference stack defines the coordinate system used for drift estimation throughout cyclic imaging.

Before each imaging cycle, a single bright-field image is captured and registered against the reference stack to estimate the accumulated 3D displacement. Registration is first performed via fast Fourier transform (FFT)–based cross-correlation between the target bright-field image and the reference stack:

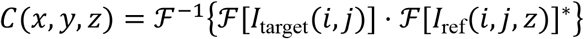

where ℱand ℱ^−1^ denote the Fourier and inverse Fourier transform, respectively. The peak of the resulting correlation volume identifies the pixel level displacement vector.

Thus, 3D displacement is refined by identifying subpixel peak positions using a phasor-based localization algorithm^26,27^, which encodes spatial position in the phase of the first Fourier coefficients:

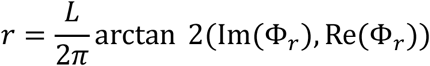

where *L* denotes the size of the extracted region of interest (11×11×11 pixels). This computation yields the 3D displacement vector Δ**r** = (Δ*x*, Δ*y*, Δ*z*) with nanometer precision (<2 s on a regular personal computer).

During STORM image acquisition, active drift correction was performed every 1000 frames (∼30 seconds). Because short-term drift is typically limited to tens of nanometers, the registration search was restricted to a 1 μm axial range to accelerate computation.

After STORM image reconstruction^37,38^, residual drift is further corrected using the AIM algorithm^28^. Localization coordinates were grouped in 100-frame intervals (∼ 3 seconds), and AIM algorithm was performed to remove remaining high-frequency drift in the final reconstructed image.

### Cell culture, fixation and immunofluorescence staining

NIH-3T3 cell lines were maintained in Dulbecco’s Modified Eagle Medium (DMEM; Gibco 10567022) supplemented with 10% fetal bovine serum (FBS; Gibco A5256801) and 1% penicillin-streptomycin (Pen/Strep; Gibco 15140163). Cells were cultured in a humidified incubator at 37 °C with 5% CO_2_. Prior to imaging, cells were seeded onto gelatin-coated ibidi µ-Slide I chambers (ibidi, cat. no. 80176) and then fixed with 4% paraformaldehyde (PFA; Electron Microscopy Sciences, cat. no. 15710) in phosphate-buffered saline (PBS, pH 7.2; Life Technologies, cat. no. 20012-019) for 10 minutes at room temperature. Following fixation, cells were washed three times with PBS. Residual aldehydes were quenched using freshly prepared 0.1% NaBH_4_ (Sigma-Aldrich, cat. no. 71320) in PBS for 7 minutes. Samples were washed three additional times with PBS (5 minutes each) prior to permeabilization using 0.2% (v/v) Triton X-100 (Sigma-Aldrich, cat. no. T8787) in PBS for 10 minutes. Samples were subsequently blocked in PBS containing 6% bovine serum albumin (BSA; Sigma-Aldrich, cat. no. A9418) and 3% normal alpaca serum (Jackson ImmunoResearch, cat. no. 028-000-121) for 1 hour at room temperature to reduce non-specific binding. For each staining/imaging cycle, antibodies were diluted in the same blocking buffer and incubated with the cells using one of three formats (see Table 1): Alexa Fluor 647-conjugated primary antibodies (1º Ab) for 2 hours at room temperature; premixed antibody and Alexa Fluor 647-conjugated secondary nanobody complex (1º Ab + 2º Nb); or unconjugated primary antibodies followed by Alexa Fluor 647-conjugated secondary nanobody (1º Ab → 2º Nb). After incubation, samples were washed three times with PBS.

**Table 1.** Antibodies used in each staining/imaging cycle.

| Cycle | Target | Primary Antibody | Host | Dilution | Detection Strategy |
| --- | --- | --- | --- | --- | --- |
| 1 | TOM20 | Abcam 186735 | Rabbit | 1:200 | $1^\circ\text{Ab} \rightarrow 2^\circ\text{Nb}$ |
| 2 | H3K9me3 | Abcam ab8898 | Rabbit | 1:400 | $1^\circ\text{Ab} + 2^\circ\text{Nb}$ |
| 3 | α-Tubulin | BioLegend 627908 | Mouse | 1:200 | $1^\circ\text{Ab}$ |
| 4 | H3K27me3 | CST 12158S | Rabbit | 1:400 | $1^\circ\text{Ab}$ |
| 5 | RNAP2S5P | Abcam ab237277 | Rabbit | 1:200 | $1^\circ\text{Ab}$ |
| 6 | H3K4me3 | Abcam ab8580 | Rabbit | 1:400 | $1^\circ\text{Ab} + 2^\circ\text{Nb}$ |

### Immunofluorescence staining

#### Alexa Fluor 647-Conjugated Primary Antibody (1º Ab)

Alexa Fluor 647-conjugated primary antibodies were diluted in the blocking buffer and incubated with samples for 2 hours at room temperature. After incubation, samples were washed three times with washing buffer consisting of 1% BSA and 0.05% Triton X-100, with 5 minutes for each wash prior to imaging.

#### Unconjugated Primary Antibodies followed by Secondary Nanobodies (1º Ab → 2º Nb)

Unconjugated primary antibodies were diluted in blocking buffer and incubated with samples for 2 hours at room temperature. Following antibody incubation, samples were washed three times with washing buffer consisting of 1% BSA and 0.05% Triton X-100 in PBS, with 5 minutes per wash. Alexa Fluor 647-conjugated anti-rabbit IgG nanobodies (Jackson ImmunoResearch, 611-604-215) were applied for 1 hour at room temperature. Samples were subsequently washed three times prior to imaging

#### Premixed Primary Antibodies and Secondary Nanobodies (1º Ab + 2º Nb)

To enable one-step immunofluorescence staining using primary antibodies from the same host species, we used premixed complexes of unconjugated primary antibody and Alexa Fluor 647-conjugated secondary nanobody. Antibody-nanobody complexes were generated by incubating unconjugated primary antibodies (1°Ab) with Alexa Fluor 647-conjugated secondary nanobodies (2°Nb; Jackson ImmunoResearch, 611-604-215) at a volume ratio of 1:4 overnight at 4°C. To block the residual Fc-binding sites, excess unlabeled nanobodies (Jackson ImmunoResearch, cat. no. 611-004-215) were added to a final concentration of 10 µg/mL and incubated for 15 minutes at room temperature prior to the application to the sample. The premixed complexes were diluted in blocking buffer and applied to samples for 2 hours at room temperature, followed by three washing steps before imaging.

### Preparation of Nitrogen-purged STORM Imaging Buffer

STORM imaging buffer consisted of 10% (w/v) glucose (Sigma-Aldrich, G8270), 0.56 mg/mL glucose oxidase (Sigma-Aldrich, G2133), 0.17 mg/mL catalase (Sigma-Aldrich, C40) and 0.14 M β-mercaptoethanol (Sigma-Aldrich, M6250). Prior to buffer preparation, the individual components of the STORM imaging buffer were first equilibrated inside a nitrogen-purged glove box for at least 30 minutes before mixing. All reagents were then combined inside the nitrogen-purged glove box to minimize oxygen exposure. The prepared imaging buffer was subsequently transferred to a nitrogen-purged incubator for storage. When maintained in a nitrogen environment, the imaging buffer can retain stable performance for up to one week.

### Fluorophore Inactivation

To prevent signal carryover between imaging cycles, residual fluorophores were chemically inactivated using meta-chloroperoxybenzoic acid (mCPBA, Sigma-Aldrich, 273031). The m-CPBA stock was prepared by dissolving m-CPBA powder in 100% ethanol at 1M concentration or 172mg/mL. Following completion of STORM image acquisition, samples were washed three times with PBS to remove imaging buffer components. A freshly prepared 10 mM mCPBA solution (diluted 1:100 from stock) was then introduced to the sample via fluidic exchange and incubated with the sample for 5 minutes at room temperature to inactivate Alexa Fluor 647. After the initial inactivation step, the samples were washed five times with PBS (1-2 minutes per wash) to remove residual oxidizing agent. A second 5 minutes of incubation with 10 mM mCPBA was subsequently performed to ensure complete fluorophore inactivation. Following the second bleaching step, samples were thoroughly rinsed with PBS to remove remaining mCPBA prior to the next staining cycle.

## Acknowledgements

This work was supported by the National Institute of Health grants R01CA301488, R01CA254112, and P41EB031772, as well as Cancer Center of Illinois Seed Grant from the University of Illinois Urbana-Champaign.

## Author Contributions

**H.M**. conceived the study; engineered the super-resolution imaging platform and the AutoStainer fluidic system; developed the bright-field-based active 3D drift correction and registration algorithms; established the mCPBA chemical quenching and the nitrogen-purged strategy; performed data analysis; drafted the manuscript and provided project supervision.

**C.Z**. developed the Python-integrated control framework for automated staining and image acquisition and performed the technical validation of the nitrogen-purged imaging buffer.

**S.Z**. prepared biological specimens and conducted cyclic (d)STORM imaging experiments to evaluate mCPBA quenching performance across multiple cycles.

**S.C**. contributed to the mechanical implementation, fluidic integration, and hardware setup of the AutoStainer module.

**Y.L**. provided critical resources, administrative and scientific support, drafted the manuscript and supervised the project.

All authors contributed to the final editing and approved the manuscript for submission.

